# Mapping Interactome Networks of FOSL1 and FOSL2 in Human Th17 Cells

**DOI:** 10.1101/2021.05.12.443731

**Authors:** Ankitha Shetty, Santosh D. Bhosale, Subhash Kumar Tripathi, Tanja Buchacher, Rahul Biradar, Omid Rasool, Robert Moulder, Sanjeev Galande, Riitta Lahesmaa

**Affiliations:** Turku Bioscience Centre, University of Turku and Åbo Akademi University; Turku 20520, Finland; InFLAMES Research Flagship Center, University of Turku; Centre of Excellence in Epigenetics, Department of Biology, Indian Institute of Science Education and Research (IISER); Pune 411008, India; Department of Biochemistry and Molecular Biology, Protein Research Group, University of Southern Denmark; Campusvej 55, Odense M, DK-5230, Denmark; The Leibniz Institute for Natural Product Research and Infection Biology – Hans Knöll Institute; Beutenbergstraße 13, 07745 Jena, Germany

**Keywords:** FOSL1, FOSL2, AP-1 transcription factors, Interactome, Immunoprecipitation, Affinity purification mass-spectrometry, Shared interactors, Parallel Reaction Monitoring (PRM)

## Abstract

Dysregulated function of Th17 cells has implications in immunodeficiencies and autoimmune disorders. Th17 cell-differentiation is orchestrated by a complex network of transcription factors, including several members of the activator protein (AP-1) family. Among these, FOSL1 and FOSL2 influence the effector responses of Th17 cells. However, the molecular mechanisms underlying these functions are unclear, owing to the poorly characterized protein interaction networks of these factors. Here, we establish the first interactomes of FOSL1 and FOSL2 in human Th17 cells, using affinity purification–mass spectrometry analysis. In addition to the known JUN proteins, we identified several novel binding partners of FOSL1 and FOSL2. Gene ontology analysis found a major fraction of these interactors to be associated with RNA binding activity, which suggests new mechanistic links. Intriguingly, 29 proteins were found to share interactions with FOSL1 and FOSL2, and these included key regulators of Th17-fate. We further validated the binding partners identified in this study by using parallel reaction monitoring targeted mass-spectrometry and other methods. Our study provides key insights into the interaction-based signaling mechanisms of FOSL factors that potentially govern Th17 cell-differentiation and associated pathologies.

## INTRODUCTION

Th17 cells are pro-inflammatory players that protect mucosal surfaces from extracellular pathogens. They can be derived *in vitro* by activating naive CD4^+^ T cells in the presence of IL-6, TGF-β and IL-1β (or IL-23). These cells are mainly characterised by the expression of IL-17A and IL17F; however, they also secrete other cytokines, such as IL-21, IL-22 and GM-CSF ^1-6^. Deficiency of Th17 cells causes susceptibility to mucocutaneous candidiasis ^7^, whereas their uncontrolled activity can result in autoimmune conditions such as rheumatoid arthritis, multiple sclerosis and systemic lupus erythematosus ^8^. To investigate the incidence of these associated diseases and design suitable therapeutic measures, it is crucial to first understand the molecular basis of Th17 cell-function.

Th17 cell-differentiation is initiated by the coordinated action of early expressed transcription factors (TFs), such as BATF, STAT3 and IRF4 ^9^. This process is also modulated by members of the activator protein (AP-1) family, which includes the JUN (JUNB ^10,11^), FOS (FOSL1 ^12^, FOSL2 ^9^) and ATF (BATF ^13^, ATF3 ^14^) TFs. FOSL1 and FOSL2 (collectively termed FOS-like proteins) are two paralogous factors that regulate embryonic development, cancer progression and immune cell signaling ^15-18^. Their significance in initiating murine Th17 responses, however, was only recently realized ^9,12^. Though molecular networks are highly conserved in human and mouse, genetic studies across the two species have revealed striking discrepancies ^5,19-23^. In light of this, a parallel study from our laboratory used cord blood T cells to verify the human-specific roles of FOSL1 and FOSL2 in Th17-regulation ^24^. Functional genomics approaches revealed that both factors negatively influence Th17 responses in human ^24^. Nonetheless, the molecular mechanisms that mediate these effects are not understood.

AP-1 TFs tend to bind to similar genomic sequences ^9,10,25^, but perform substantially different functions ^18^. Such versatility might be achieved through a dynamic interactome ^26-28^. FOSL1 and FOSL2 lack a transactivation domain and, thus, must interact with JUN and other proteins to regulate their target genes. Furthermore, since these factors occupy DNA as a dimer, their regulatory abilities are significantly influenced by their interacting partners ^28^. Despite extensive research on AP-1 signaling, the global interactomes of AP-1 TFs are largely unexplored in T cells. Mapping the interaction networks of FOSL1 and FOSL2 in human Th17 cells can thus advance our understanding of their signaling mechanisms in this milieu.

Affinity purification–mass spectroscopy (AP-MS) has emerged as a reliable method for identifying protein-protein interactions (PPIs) at a global level ^29-31^. MS in particular, detects and quantifies proteins in an unbiased manner, without prior knowledge. In the present study, we co-immunoprecipitated putative interactors of FOSL1 and FOSL2 in human Th17 cells, and identified them by liquid chromatography–tandem MS (LC-MS/MS). Our analysis is the first to compare the FOSL1 and FOSL2 interactomes, thereby revealing their shared and unique binding partners. Parallel reaction monitoring targeted MS (PRM-MS) and immunoblotting (IB) were used to reliably validate the top interactors of these factors. Together with the predicted functionalities of the FOSL PPI networks, this study delivers a perspective on how FOSL proteins could regulate human Th17 cell-identity. Such comprehensive analysis could help gain crucial insights into new therapeutic strategies for treating autoimmune diseases.

## METHODS

### Human CD4+ T-cell isolation

Mononuclear cells were isolated from the human umbilical cord blood of healthy neonates (Turku University Central Hospital, Turku, Finland) by Ficoll-Paque density gradient centrifugation (Ficoll-Paque PLUS; GE Healthcare). Naive CD4^+^ T cells were further purified using CD4+ Dynal positive selection beads (Dynal CD4 Positive Isolation Kit; Invitrogen), by following the manufacturer’s protocol.

### *In vitro* culturing of Th17 cells

CD4+ T cells were activated with plate-bound α-CD3 (3.75 µg/ml; Immunotech) and soluble α-CD28 (1 μg/mL; Immunotech) in X-vivo 20 serum-free medium (Lonza). X-vivo 20 medium was supplemented with L-glutamine (2 mM, Sigma-Aldrich) and antibiotics (50 U/mL penicillin and 50 μg/mL streptomycin; Sigma-Aldrich). Th17-cell differentiation was induced using a cytokine cocktail of IL-6 (20 ng/mL; Roche), IL-1β (10 ng/mL) and TGF-β (10 ng/mL) in the presence of the neutralizing antibodies anti-IFN-γ (1 μg/mL) and anti-IL-4 (1 μg/mL) to block Th1 and Th2 polarization, respectively. For the control cells (Th0), CD4+ T cells were TCR stimulated with α-CD3 and α-CD28 in the presence of neutralizing antibodies. All cytokines and neutralizing antibodies were purchased from R&D Systems, unless otherwise stated. All cultures were maintained at 37°C in a humidified atmosphere of 5% (v/v) CO_2_/air.

### IL-17 secretion

Secreted IL-17A levels were estimated in cell-culture supernatants of 72h Th17 cells using human IL-17A Duoset ELISA kit (R&D Biosystems DY317-05, DY008). The amount of IL-17A secreted by Th17 cells was normalized with the number of living cells, based on forward and side scattering in flow cytometry analysis (LSRII flow cytometer; BD Biosciences).

### Flow cytometry

Th17 cells were harvested at 72h, washed with FACS buffer (0.5% FBS; 0.1% Na-azide; PBS), and further incubated with PE-labelled anti-CCR6 antibody (BD Cat no. 559562) for 20 min at 4°C. Suitable isotype controls were used. Samples were analysed using LSRII flow cytometer (BD Biosciences). Live cells were gated based on forward and side scattering. The acquired data were analysed with FlowJo (FLOWJO, LLC).

### Western blotting

Cell culture pellets were lysed using RIPA buffer (Pierce, Cat no. 89901) that was supplemented with protease and phosphatase inhibitors (Roche) and sonicated using a Bioruptor UCD-200 (Diagenode). Sonicated lysates were centrifuged at 14,000 rpm for 30 min at 4°C, and supernatants were collected. Samples were estimated for protein concentration (DC Protein Assay; Bio-Rad) and boiled in 6x Laemmli buffer (330 mM Tris-HCl, pH 6.8; 330 mM SDS; 6% β-ME; 170 μM bromophenol blue; 30% glycerol). Samples were then loaded on gradient Mini-PROTEAN TGX Precast Protein Gels (BioRad) and transferred to PVDF membranes (Trans-Blot Turbo Transfer Packs, BioRad).

For protein expression analysis of FOSL1 and FOSL2, the following antibodies were used: anti-FOSL1 (Cell Signaling Tech, Cat no. 5281), anti-FOSL2 (Cell Signaling Tech., Cat no.19967) and anti-β-actin (SIGMA, Cat no. A5441). HRP-conjugated anti-mouse IgG (SantaCruz, Cat no. sc-2005) and anti-rabbit IgG (BD Pharmingen, Cat no. 554021) were used as secondary antibodies.

### Cellular fractionation

Cell pellets of Th0 and Th17 cultures (24h and 72h) were lysed and fractionated into cytoplasmic and nuclear components using the NE-PER Nuclear and Cytoplasmic Extraction Reagent Kit, (Thermo Fischer Scientific, Cat no. 78833), by following the manufacturer’s instructions. Extracts were then analysed by western blotting. FOSL localization was determined using anti-FOSL1 (Cell Signaling Tech, Cat no. 5281) and anti-FOSL2 (Cell Signaling Tech., Cat no.19967) antibodies. Anti-GAPDH (Hytest, Cat no. 5G4) and anti-LSD1 (Diagenode, Cat no. C15410067) antibodies were used to mark the cytoplasmic and nuclear fractions, respectively.

### Immunoprecipitation

Immunoprecipitation (IP) for FOSL1 and FOSL2 was performed using Pierce MS-Compatible Magnetic IP Kit (Thermo Fischer, Cat no.90409). Cell pellets from 72h cultured Th17 cells were lysed in appropriate volumes of lysis buffer provided in the kit. FOSL1 (Santacruz Biotechnology, Cat no.sc-28310), FOSL2 (Cell Signaling Technology, Cat no.19967), mouse IgG (negative control for FOSL1; Cell Signaling, Cat no. 5415), and rabbit IgG (negative control for FOSL2; Cell Signaling Technology, Cat no. 2729) antibodies were used for IP of the respective protein complexes. All antibodies were pre-incubated with protein A/G beads for 4–5 h to form antibody-bead complexes. Lysates were first pre-cleared with control IgG-bead complexes, for 3 h. Pre-cleared lysates were then incubated overnight with FOSL1/FOSL2 antibody-bead complexes (test IP) or the corresponding control IgG-bead complexes (negative IP control). The pull-down fractions were washed (following the manufacturer’s protocol) and eluted with appropriate volume of elution buffer. The eluted protein was vacuum dried for MS analysis or run for western blotting.

The antibodies used for IP-Immunoblotting are as follows: anti-FOSL1 (Santacruz Biotechnology, Cat no.sc-28310), anti-FOSL2 (Cell Signaling Technology, Cat no.19967), anti-RUNX1 A-2 (Santa Cruz Biotechnology, Cat no. sc-365644); anti-JUNB C-11 (Santa Cruz Biotechnology, Cat no.sc-8051); anti-SIRT1 (Cell Signaling Technology, Cat no. 2496); anti-JUN (BD Biosciences, Cat no.610326). Conformation-specific rabbit HRP (Cell Signaling Technology, Cat no.5127) and mouse HRP (Cell Signaling Technology, Cat no. 58802) were used as secondary antibodies.

### Sample preparation for mass-spectrometry analysis

The IP eluates for control IgG, FOSL1 and FOSL2 were denatured with urea buffer (8 M urea, 50 mM Tris-HCl, pH 8.0), followed by reduction using dithiothreitol (10 mM) at 37°C for 1 h. The reduced cysteine residues were subsequently alkylated using iodoacetamide (14 mM, in darkness) at room temperature for 30 min. The samples were diluted to reduce the urea concentration (< 1 molar), followed by digestion with sequencing grade modified trypsin at 37°C overnight (16–18 h). The digested peptides were acidified and then desalted using C18 Stage Tips, prepared in-house using Empore C18 disks (3M, Cat. no 2215). The desalted samples were dried in a SpeedVac (SAVANT SPD1010, Thermo Scientific) and then stored at -80°C until further analysis.

For validation measurements, synthetic isotopic analogues (lysine ^13^C_6_ ^15^N_2_ and arginine ^13^C_6_ ^15^N_4_) were obtained for unique peptides from selected protein targets identified in the AP-MS discovery data (Thermo Fischer Scientific). The same sample preparation procedure was used for the validation experiments, with the exception that the samples were spiked with isotope-labelled peptides and MSRT retention time peptides standards (Sigma), prior to MS analysis.

### LC-MS/MS Analysis

#### A. Data-Dependent Analysis

The dried peptides were reconstituted in formic acid/acetonitrile (both 2% in water), and a NanoDrop-1000 UV spectrophotometer (Thermo Scientific) was used to measure the peptide amounts. Equivalent aliquots of the digested peptides were analysed by LC-MS/MS using an Easy-nLC 1200 coupled to Q Exactive HF mass spectrometer (Thermo Fisher Scientific). The peptides were loaded onto a 20 × 0.1 mm i.d. pre-column and separated with a 75 µm x 150 mm analytical column, both packed with 5 µm Reprosil C18 (Dr Maisch GmbH). A separation gradient of 5–36% B in 50 min was used at a flow rate of 300 nl/min (Solvent A: 0.1% formic acid in MiliQ H_2_O and Solvent B: 80% acetonitrile, 0.1% formic acid in MiliQ H_2_O). The tandem MS spectra were acquired in positive ion mode with a data-dependent Top 15 acquisition method at 300–1750 m/z using HCD fragmentation. The singly and unassigned charged species were excluded from the fragmentation. MS1 and MS/MS spectra were acquired in the Orbitrap, at a resolution set to 120,000 and 15,000 (at m/z 200), respectively. The AGC target values for MS1 and MS/MS were set to 3,000,000 and 50,000 ions, with maximal injection times of 100 and 150 ms, respectively, and the lowest mass was fixed at m/z 120. Dynamic exclusion was set to 20 s. Triplicate analysis were performed for all samples in randomized batches.

#### B. Parallel reaction monitoring

Synthetic peptide analogues for validation targets were analysed together with MSRT retention time peptides standards (Sigma) by LC-MS/MS using an Orbitrap Fusion Lumos mass spectrometer, coupled to Easy-nanoLC (Thermo Fisher Scientific) with the same column configuration as above. On the basis of these data, a PRM method was developed for the analysis of these targets and their endogenous counterparts in AP validation samples. For the targeted analysis, the peptides were separated with a 30-min gradient of 8–39% solvent B. Data were acquired in a PRM mode with an isolation window setting of 1.6 m/z at a resolution of 15,000 for the Orbitrap, using a target AGC value of 50,000 and maximum injection time of 22 ms.

### Data Analysis

#### A. AP-MS Data

The MS raw files were searched against a UniProt FASTA sequence database of the human proteome (downloaded, May 2019, 20415 entries:) using the Andromeda search engine, incorporated with the MaxQuant software (Version 1.6.0.16) ^32,33^. Trypsin digestion, with amaximum of two missed cleavages, carbamidomethylation of cysteine as a fixed modification, and variable modification of methionine oxidation and N-terminal acetylation were specified in the searches. A false discovery rate (FDR) of 1% was applied at the peptide and protein levels. MaxQuant’s label-free quantification (LFQ) algorithm ^34^ was used to calculate the relative protein intensity profiles across the samples. The “match between run” option was enabled to perform matching across the MS measurements.

The proteinGroup.txt file from the MaxQuant output was further processed using Perseus (Version 1.6.2.3) ^35^. The output was filtered to remove contaminants, reverse hits and proteins only identified by site. Protein LFQ values were log2 transformed, and the medians of the technical replicates calculated. The data were filtered to retain proteins with three valid values in at least one group (IgG, FOSL1 and FOSL2 pulldown). The resulting data matrix was then analysed using the MS interaction statistics (MiST) algorithm. The algorithm calculates a MiST score for each of the potential interactors on the basis of their intensity, consistency and specificity to the bait ^36^. MiST score criteria of ≥ 0.75 for FOSL1 and FOSL2-prey interaction and ≤ 0.75 for interaction with IgG were applied. Further, to eliminate proteins frequently detected as contaminants in IP experiments, comparison was made with a list of proteins that were detected with IgG-mock baits in other T-helper cell studies of our laboratory (these were based on 126 other IP experiments). We retained those proteins that were detected with a frequency of less than 40% in the described in-house database for possible contaminants. Finally, we listed the top binding partners of FOSL1 and FOSL2 based on their abundance values in respective FOSL IP, as compared to the corresponding IgG control.

Heatmaps for the subsequent list of FOSL1 and FOSL2 interactors, identified across three biological replicates (Pulldown vs. IgG), were plotted using the Perseus software. The grey colour in the heatmaps represent undetected proteins in the respective IP experiments. The interactors were additionally mapped against the STRING database, and the assigned PPI networks were further visualized using Cytoscape ^37^.

#### B. Validation Data

Data from analysis of the synthetic peptides were analysed using Proteome Discoverer (Version 2.2, Thermo Fisher Scientific) and a FASTA file containing the sequences of the peptide targets. The MSF file from Proteome Discoverer was then used to construct spectral library in Skyline (v4.2) software ^38^ and define their retention time indices. Skyline was then used to create scheduled isolation lists for PRM analysis ^38^, process the PRM-MS raw files, and review the transitions and integration of the peptide peaks. The transition signals of endogenous peptides were normalized to their heavy counterparts, and the statistical analysis was performed using in built MSStat plugin ^39^ on the basis of sum of transition areas.

#### C. Data availability

The mass spectrometry proteomics data have been deposited to the ProteomeXchange Consortium via the PRIDE ^40^ partner repository with the dataset identifier PXD025729. The details of PRM-mass spectrometry measurements can be found in the Skyline Panorama ^41^ link https://panoramaweb.org/FOSL1_2_Th17.url and are deposited to the ProteomeXchange Consortium via the PRIDE partner repository with the data set identifier PXD025840. Reviewer Account details:

1) **Project accession:** PXD025729

Username

Password: vp0sPjNd

2)**Project accession:** PXD025840

Password: jxPwigJO

### Cellular component analysis using Ingenuity Pathway Analysis

To map the cellular locations of the identified interactors, the list of binding partners was annotated using Ingenuity Pathway Analysis (IPA, www.qiagen.com/ingenuity; Qiagen; March 2019) tool.

### GO functional enrichment analysis and networks

GO molecular function pie charts and networks were created using ClueGO and CluePedia plugins from Cytoscape, based on the p value ≤ 0.05 and corrected using a Bonferroni step-down method.

### Graphical representation, Venn diagrams and Statistical Analysis

All graphs were plotted using GraphPad Prism software (V8.3.0). Two-tailed students T-test was used to calculate statistical significance. Venn diagrams were generated using Biovenn^42^.

### Graphical illustration for workflow of the study

The pictorial representation for workflow in Fig. 1B was created using BioRender.com.

**Fig. 1.**
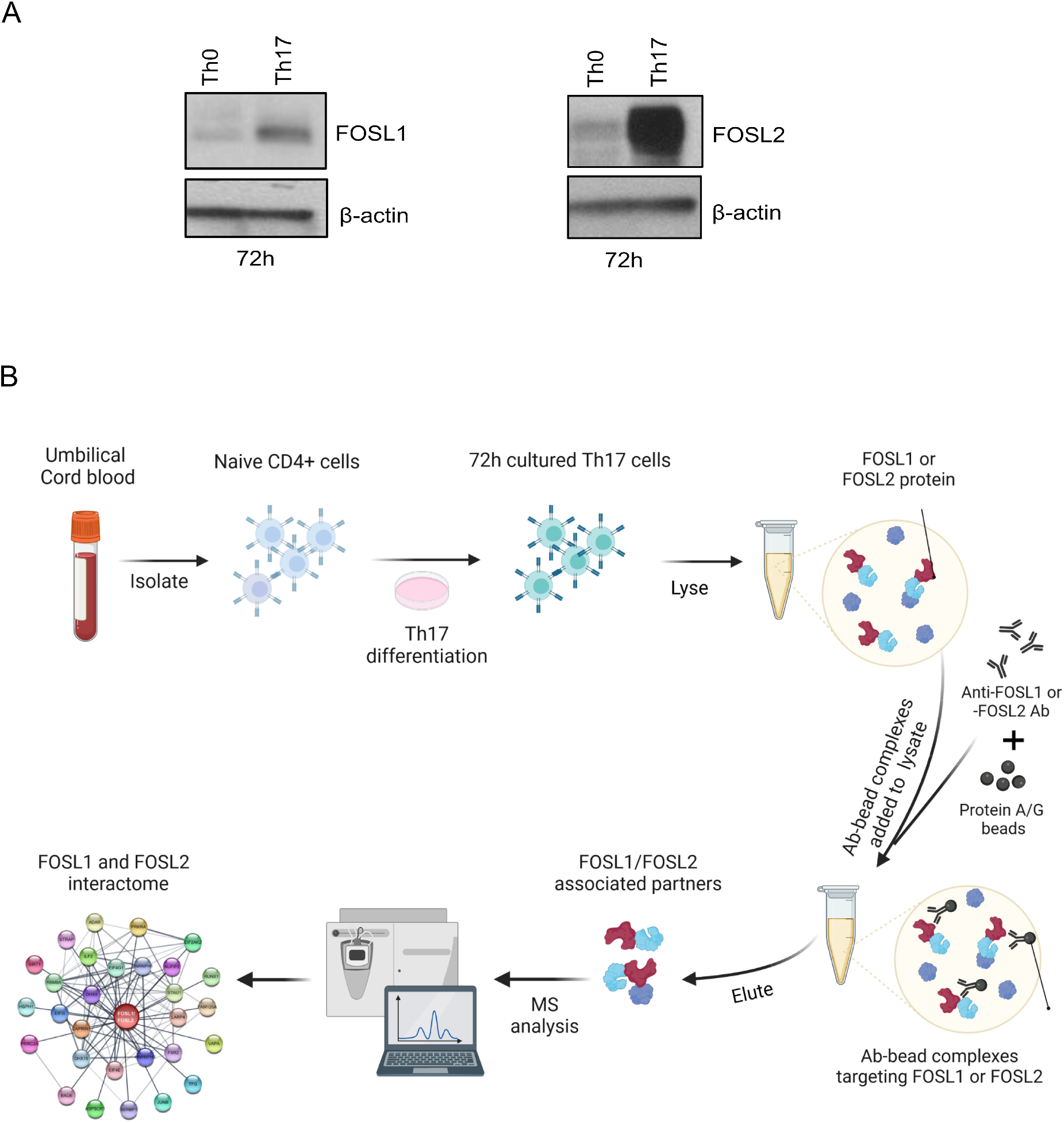
FOSL1 and FOSL2 expression and workflow for their proteomic analysis in human Th17 cells. **A**. Immunoblots show expression of FOSL1 (left) and FOSL2 (right) in naive CD4^+^ T cells cultured under activating (Th0) or Th17-polarizing conditions for 72h. Actin was used as loading control. Blots for one of the three biological replicates are shown. **B**. Workflow for the study. Naive CD4^+^ T cells were isolated from human umbilical cord blood and polarized to Th17 phenotype for 72h. The cultured cells were lysed, and FOSL1 or FOSL1 protein was immunoprecipitated using their respective antibodies. The pull-down fractions were then analyzed for binding partners of FOSL1 or FOSL2 using LC-MS/MS based protein interactome analysis.

## RESULTS

### FOS-like proteins are upregulated during initiation of human Th17 cell differentiation

To study the FOSL1 and FOSL2 interactomes in early-differentiating human Th17 cells, we first determined their expression in naive CD4^+^ T cells that were *in vitro* stimulated and polarized toward Th17-fate for 72h. The polarization efficiency was confirmed, based on the consistent expression of the lineage-defining markers, CCR6 and IL-17 cytokine (Fig. S1A and B). Immunoblot analysis of the 72h cultures revealed a Th17-specific increase in levels of FOSL1 and FOSL2 (relative to activated (Th0) cells), which made it the preferred time point for our proteomic analysis (Fig. 1A, Fig. S1C). The expression of these factors in human is consistent with previous findings in mouse ^9,12^.

### Systematic analysis unravels FOSL1 and FOSL2 interacting partners in human Th17 cells

To identify the interacting partners of FOSL1 and FOSL2, an AP-MS approach was used. Workflow for the current study is illustrated in Fig. 1B. Th17 cells from three independent biological replicates were lysed, and IP was performed using FOSL1 or FOSL2 antibodies (Ab), as well as the corresponding species-specific control IgG antibodies. Pull-down of the bait proteins (FOSL1 or FOSL2) was confirmed with immunoblotting (Fig. 2A and B), and the IP fractions were then analyzed for interacting partners with LC-MS/MS. Using the MaxQuant label-free quantitation (LFQ) algorithm, the relative protein intensities were compared across the samples. The putative interactions were further prioritized by intensity, reproducibility and specificity to the bait, with the MiST algorithm ^36^. Our analysis reliably identified 163 and 67 binding partners of FOSL1 and FOSL2, respectively. These were obtained after strategically eliminating the non-specific interactions, based on (1) comparing the enrichment scores with IgG control and (2) using an in-house repository of possible contaminants derived from other AP-MS experiments in our laboratory.

**Fig. 2.**
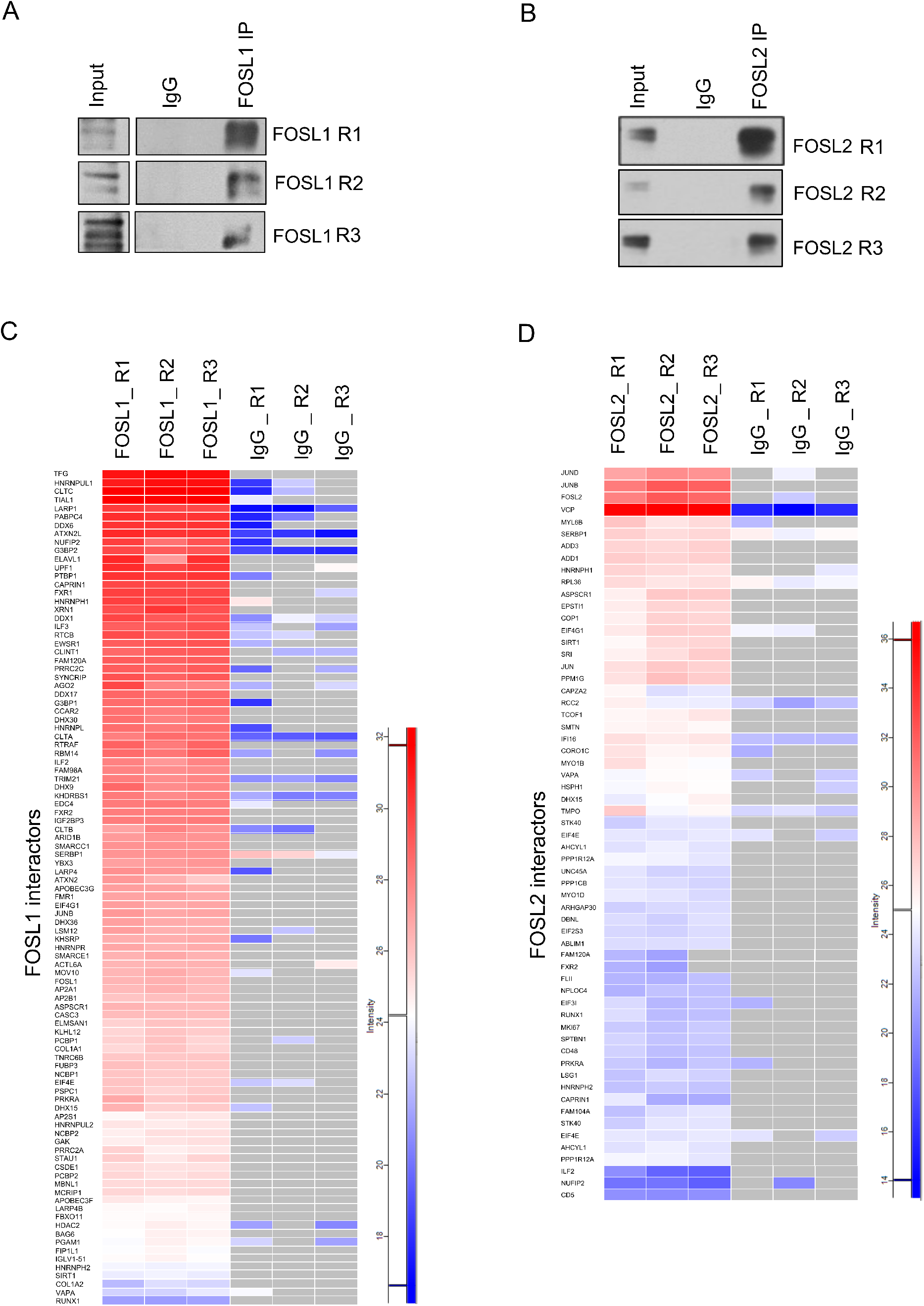
Analysis of FOSL1 and FOSL2 protein-protein interactions in human Th17 cells. **A & B**. Immunoblots confirm immunoprecipitation of FOSL1 (panel A) and FOSL2 (panel B) protein from 72h cultured Th17-cell lysates. Blots show lanes for total lysate (input), IgG control IP and FOSL1/FOSL2 IP. **C & D**. FOSL1 or FOSL2 pull-down fractions from three biological replicates (R1, R2 and R3) were analyzed for their corresponding binding partners using LC-MS/MS. Heatmaps depict Log_2_intensity values for the topmost interacting partners of FOSL1 (panel C) and FOSL2 (panel D) in 72h human Th17 cells. Grey color indicates missing or undetected proteins.

The top binding partners of FOSL1 and FOSL2 and their corresponding enrichment scores are depicted in the heatmaps of Fig. 2C and D (For all identified partners, see Fig. S2A and B). FOS-JUN dimers are one of the most widely occurring protein-protein associations, across cell-types. Amid members of the JUN family, JUNB was among the top interactors of both FOSL proteins, which agrees with previous findings ^10,12,43^. Additionally, several new binding partners were identified. These included XRN1, AP2A1, PCBP1, ILF3, TRIM21, HNRNPH1, and HDAC2 for FOSL1, and ADD3, PPP1CB, MYO1B, HNRNPH1, CD48, and CD5 for FOSL2. Intriguingly, despite being paralogs with similar functions, FOSL1 and FOSL2 showed no interaction with each other. Although their association is reported in lower organisms, such as yeast ^44^, none of the existing studies in vertebrates support such findings.

### Gene Ontology functional enrichment analysis of FOSL1 and FOSL2 interactors

Spatial organization of signaling networks relies on the cellular location of the proteins that constitute the network ^45^. FOSL proteins can be cytoplasmic or nuclear and can shuttle between these compartments, in a context-dependent manner ^46,47^. Their localization profile in human T cells, however, is yet to be studied. We addressed this by performing subcellular fractionation on Th0 and Th17 lysates (24h and 72h), which detected both proteins predominantly within nuclear fractions (Fig. 3A, Fig. S3A). To gain further insights on FOSL-mediated signaling networks, the cellular distribution of their interacting partners was determined using ingenuity pathways analysis (IPA) (Fig. 3B). More than 50% of the FOSL1 interactors and nearly 33% of the FOSL2 interactors were associated with the nucleus. Of the rest, most were cytoplasmic, and only a small fraction (10-15%) corresponded to other cellular compartments.

**Fig. 3.**
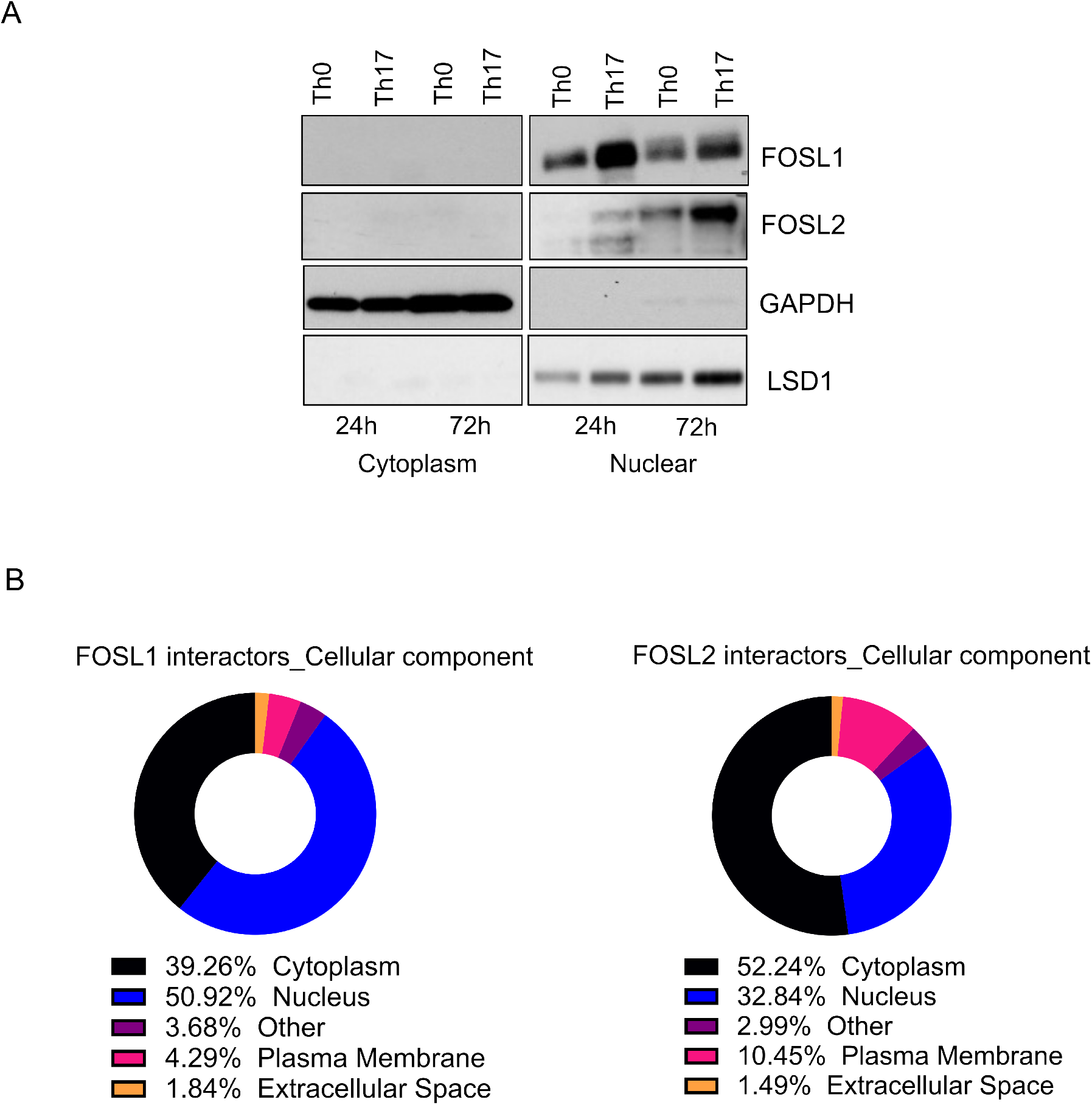
Localization of FOSL proteins and cellular distribution of their binding partners. **A**. Following subcellular fractionation of Th0 and Th17 cell lysates (24h and 72h), immunoblotting was performed to determine FOSL1 and FOSL2 expression in the fractionated samples. LSD1 and GAPDH were used to mark nuclear and cytoplasmic fractions respectively. Representative blot for three biological replicates is shown. **B**. Pie charts show classification of FOSL1 (left) and FOSL2 (right) interacting proteins based on their cellular localization, using Ingenuity Pathway Analysis (IPA).

To determine the physiological relevance of these interactions, their molecular functionalities were mapped using the GO database (Fig. S4A and B). Proteins interacting with FOSL1 were enriched for functions, such as RNA binding (60%; constituted by single-stranded, double-stranded, mRNA and siRNA binding), nucleosomal DNA binding (6.67%), mRNA 5′ UTR binding (6.67%), RNA 7-methylguanosine cap binding (16.67%), clathrin binding (6.67%), and RNA helicase activity (3.33%) (Fig. S4A). Similarly, FOSL2 interactors were enriched for translational initiation activity (66.67%), double-stranded RNA binding (16.67%) and actin filament binding (16.67%) (Fig. S4B). Remarkably, RNA binding and translational initiation constituted more than 80% of the identified functionalities for either interactomes.

The role of RNA-binding proteins (RBPs) in regulating FOS/JUN functions has been previously studied ^48-50^. RBPs also post-transcriptionally modify cytokine mRNAs, through which they modulate T-cell development, activation and differentiation ^51-54^. For instance, the zinc finger protein Tristetraprolin (TTP) is reported to impair Th17 differentiation by destabilizing IL-17 transcripts ^55,56^. TTP targets the degradation of mRNA molecules by coordinating with exonucleases such as XRN1 ^57^. Interestingly, XRN proteins (XRN1, XRN2) were detected as a part of our FOSL1 interactome along with the RBPs, UPF1 and UPF2, which are known to trigger mRNA decay ^48,58^. Apart from IL-17, several other transcripts that code for Th17-regulatory factors, such as STAT3, IRF4, CTLA4, GM-CSF, and CCR6, are reported to be controlled by the action of RBPs ^59^. These findings imply that FOSL1 might restrain Th17-signaling via association with proteins that destabilize the lineage-specific transcripts. This warrants further investigation.

Network analysis was further performed for the enriched GO functionalities by using Cytoscape (Fig. 4A and B). Within the FOSL1 interaction network, the subclusters associated with different RNA binding functions were highly interconnected, which hints at their interdependent roles. RBPs are also involved in the regulation of translational initiation through various mechanisms ^60^. This association was evident in the GO networks for both FOSL factors (Fig. 4A and B). Eukaryotic translational initiation factors (eIFs) stabilize ribosomal pre-initiation complexes and mediate post-transcriptional gene regulation ^60,61^. Within the eIF family, we found eIF4G1, eIF4E, eIF3I and eIF2AK2 to interact with FOSL1, as well as FOSL2. Out of these, eIF4E constitutes the rate limiting step for translational initiation by binding to the m7G cap of transcripts ^61^. Interestingly, eIF4E is required for the pathogenesis of EAE in mouse ^62^. It is also found to be targeted by the miRNA-467b in order to inhibit Th17 differentiation and autoimmune-development ^63^. Thus, the binding of FOSL factors to translational initiation factors points towards a unique strategy for monitoring Th17 responses.

**Fig. 4.**
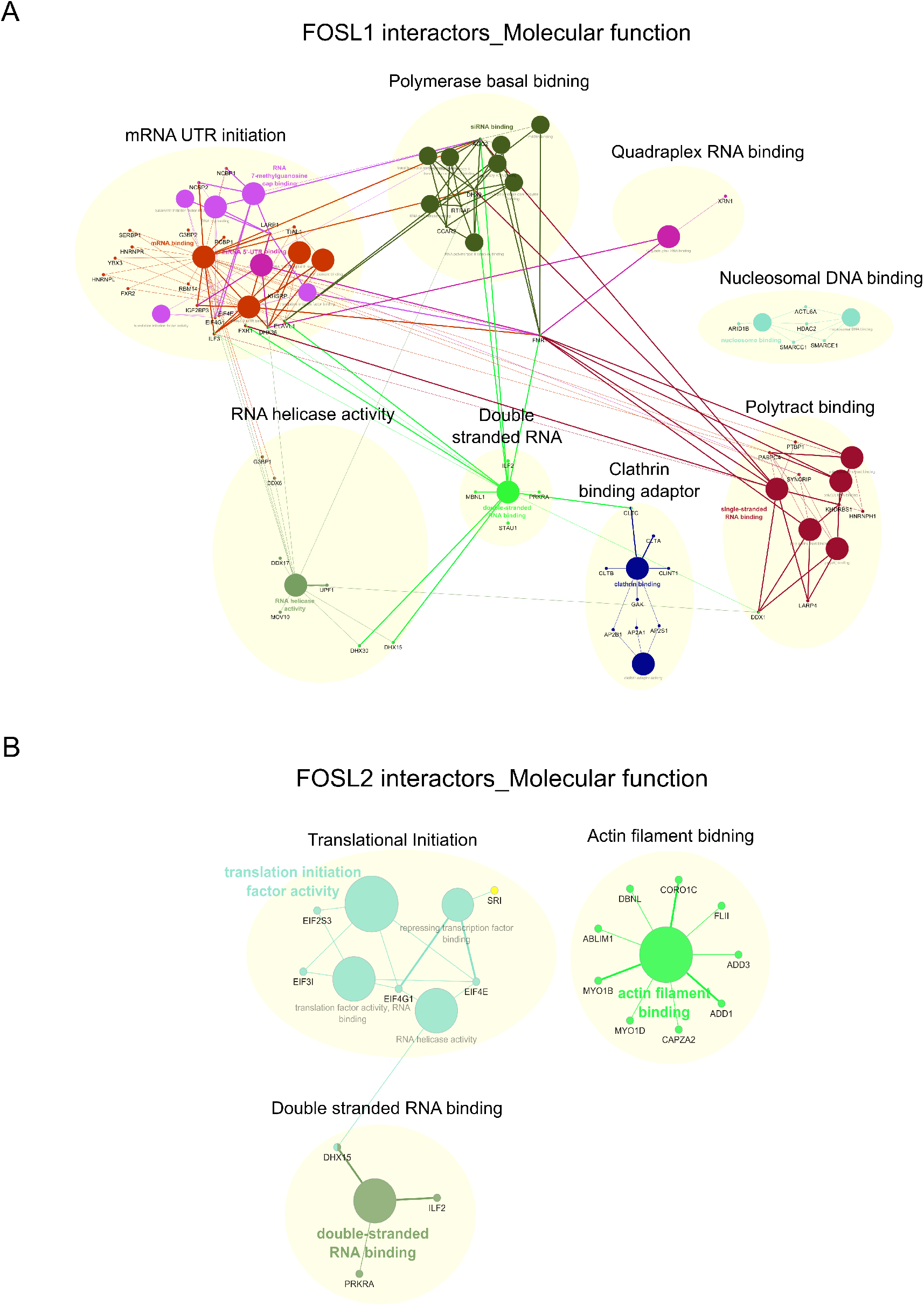
Molecular function networks enriched for FOSL1 and FOSL2 interactors. **A & B**. FOSL1 (panel A) and FOSL2 (panel B) interactors were clustered based on their molecular functions and the resulting networks were visualized using ClueGO and CluePedia plugins built in Cytoscape (Bonferroni step-down corrected p values < 0.05).

### FOSL1 and FOSL2 share interactions with key Th17 lineage-associated proteins

Prediction models have indicated that interaction partners shared by candidate TFs could facilitate co-operative or competitive tendencies between the factors ^64^. Our recent study revealed a functional coordination between FOSL1 and FOSL2 during human Th17-regulation ^24^. To investigate whether an interactome-based mechanism regulates this paradigm, we analysed these factors for their common binding partners. Our analysis revealed a total of 29 proteins to share interactions with FOSL1 and FOSL2, including JUNB, SIRT-1, HSPH1, DHX9, HNRNPH1/2, NUFIP2, LARP4, RUNX1, ADAR and EIF4E, which are associated with T-cell effector functions (Fig. 5A and B).

**Fig. 5.**
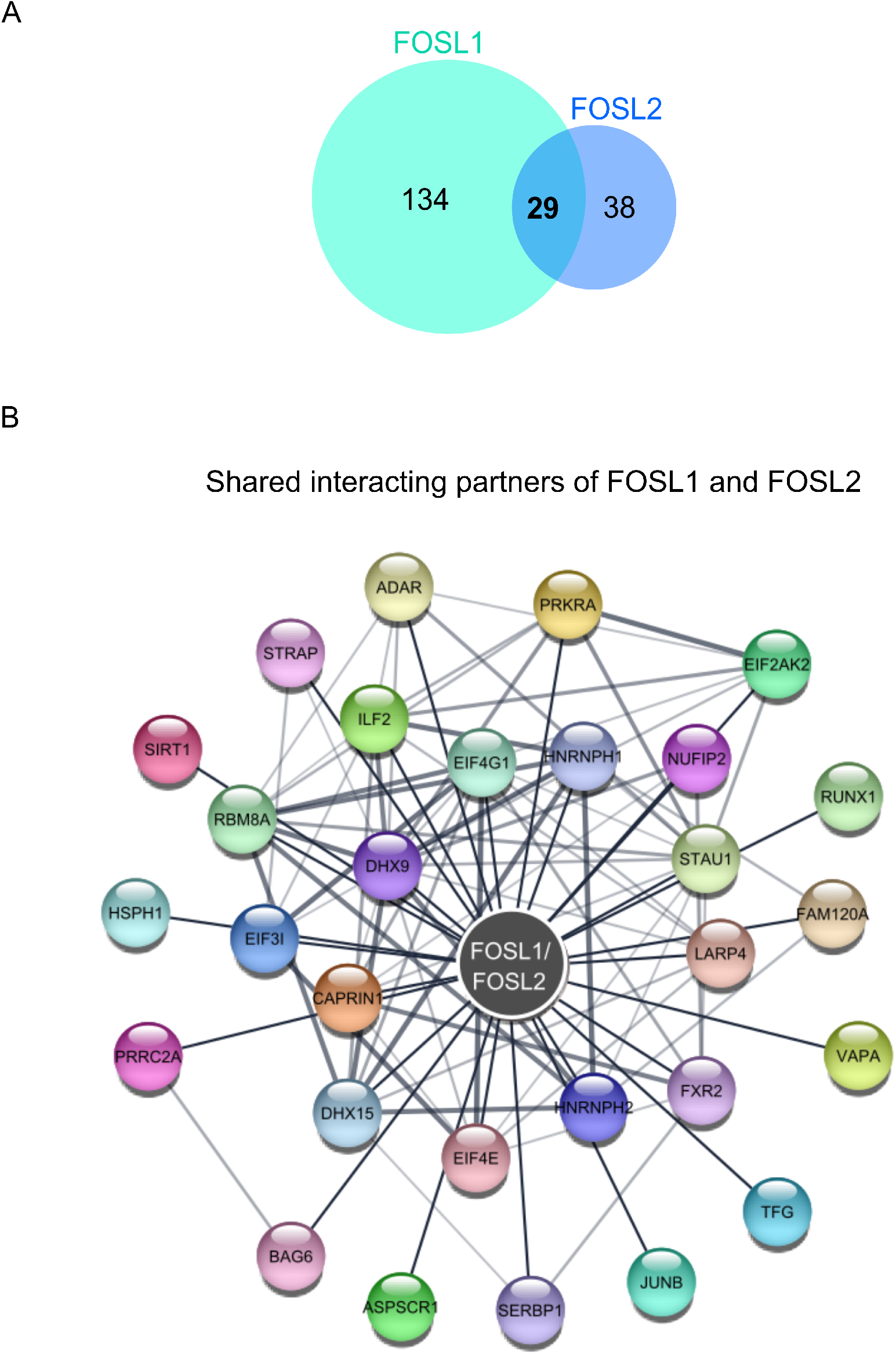
Shared interacting partners of FOSL1 and FOSL2 in human Th17 cells. **A**. Venn diagram shows the number of shared and unique interactors of FOSL1 and FOSL2 in human Th17 cells, as identified by LC-MS/MS analysis. **B**. The protein-protein interactions shared between FOSL1 and FOSL2 were mapped against the STRING database and further visualized using Cytoscape (a medium confidence score of 0.4 was used to create the STRING network)

To study these common binding partners in the context of Th17 cell-signaling, we focused on the ones that are known to regulate the lineage. These included JUNB ^10-12^, RUNX1 ^65^, SIRT-1 ^66^, eIF4E ^62,63,67^ and ADAR ^68^. Murine studies have found JUNB to promote Th17-fate and restrain alternative lineages by associating with BATF and FOSL2 ^10^. Likewise, RUNX1 and SIRT-1 are reported to influence Th17 cell-signaling. Interestingly, RUNX1’s effect on the lineage is largely governed by its binding partners. Its interaction with FOXP3 inhibits Th17-differentiation, whereas its association with RORγT activates the lineage ^65^. SIRT-1 analogously functions via RORγT, where it binds, deacetylates and enables the latter to promote Th17 cell-function ^66^. The above findings indicate that FOSL1 and FOSL2 may associate with key regulators of the lineage in order to modulate effector responses of Th17 cells.

The list of common partners also included numerous proteins with undetermined roles in Th17-regulation, including NUFIP2, HNRNPH1, HNRNPH2, DHX9, DHX15, SERBP1, and others (Fig. 5B). However, when evaluated in the context of other relevant studies, potential roles in controlling the Th17 lineage can be assigned to these factors. For instance, NUFIP2 acts as co-factor for the RNA binding protein Roquin ^69^, which is reported to inhibit Th17 differentiation ^70^. This is possibly mediated via the post-transcriptional repression of Th17-activators, such as ICOS ^69,71^, by the NUFIP2-Roquin complex. Our analysis also detected heterogenous nuclear ribonucleoproteins (hnRNPs), namely hnRNPH1 and hnRNPH2, which are involved in pre-mRNA alternative splicing. Interestingly, these proteins are closely associated with another member of the same family, hNRNPF, which reportedly interacts with FOXP3 ^72^. While FOXP3 is a master-regulator of Treg differentiation, it also inhibits Th17-signaling by antagonizing RORγt ^73,74^. Thus, our MS analysis provides a number of new interaction partners that imply novel mechanisms through which FOSL proteins alter Th17 cell fate.

### Experimental validation of the shared and unique binding partners of FOSL1 and FOSL2

To confirm their shared interactions with Th17-associated proteins, FOSL1 or FOSL2 was immunoprecipitated and immunoblotted to probe for JUNB, RUNX1, JUN and SIRT-1 (Fig. 6A and B; Fig. S5A and B). The results revealed a reproducible interaction of the FOSL factors with all of the assessed proteins. Although MS analysis of FOSL1 did not detect JUN, our IB findings indicate FOSL1-JUN complexes in Th17 cells.

**Fig. 6.**
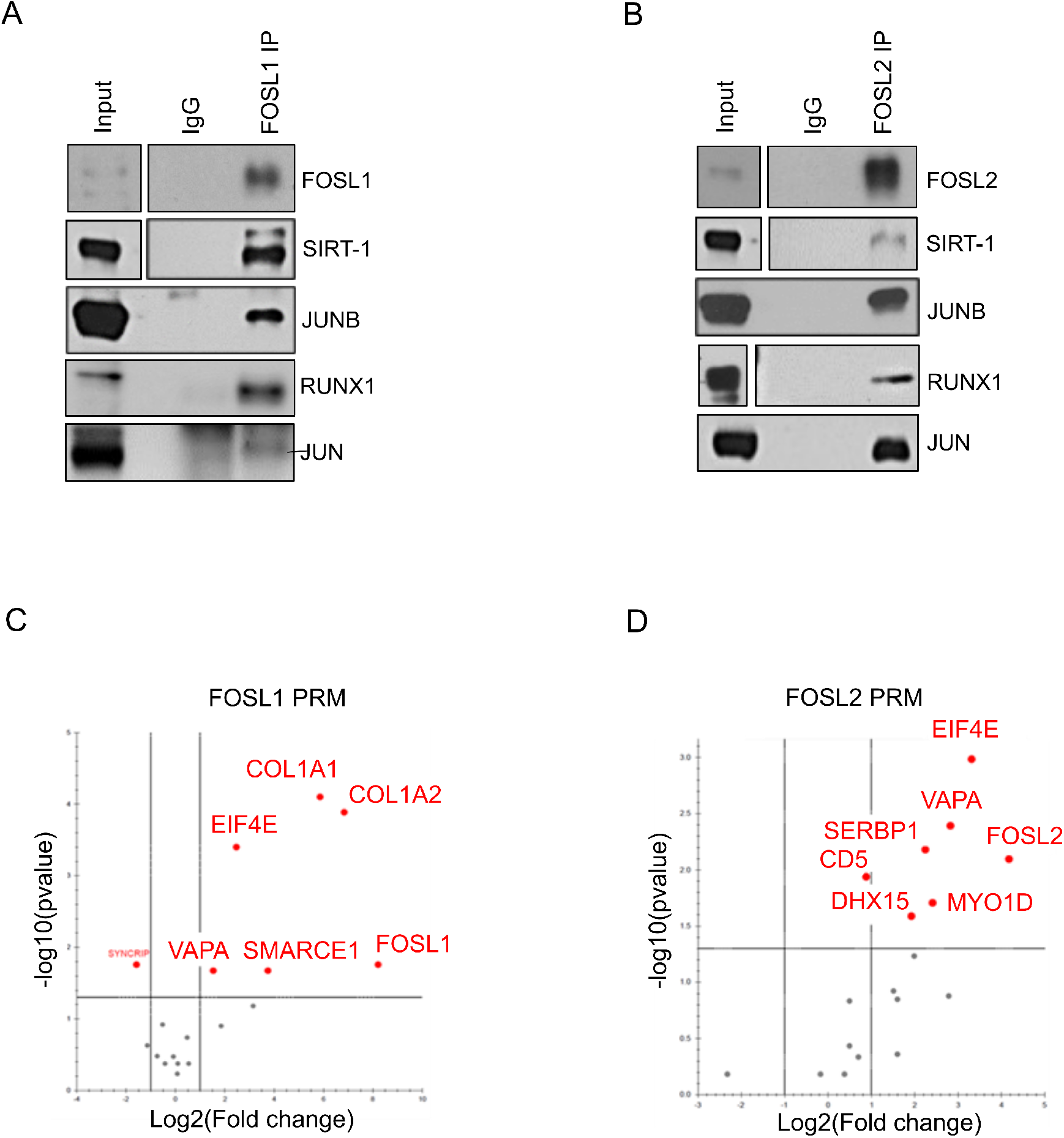
Validation of selected binding partners of FOSL1 and FOSL2. **A & B**. FOSL1 (panel A) and FOSL2 (panel B) protein was immunoprecipitated from 72h cultured Th17 cells and western blotting was used to confirm their MS-identified interactions with SIRT-1, JUNB, RUNX1 and JUN. Blots show lanes for total lysate (input), control IgG IP and FOSL1/FOSL2 IP. Figures show representative blots for two or three biological replicates (See Fig. S5 for all the replicates). **C & D**. Volcano plots show selected binding partners of FOSL1 (panel C) and FOSL2 (panel D) that were validated by Parallel Reaction Monitoring (PRM) MS analysis. Data is representative of three biological replicates. The plots were extracted using Skyline.

Even among the unique interactors, we detected several candidates that have implications in Th17-development and inflammatory phenotypes. These included the FOSL1 partners alpha-1 type I collagen (COL1A1) ^75^, SMARCE1 ^76^, TRIM21 ^77^ and HDAC2 ^78^, as well as the FOSL2 interactors MYO1D ^79^, CD48 ^80^ and JUND ^10^. A few of these, along with the other binding partners, were validated using targeted MS. With its increased sensitivity, reproducibility and ease of implementation of the technique, PRM analysis was employed to confirm selected interactors of FOSL1 (COL1A1, COL1A2, SMARCE1, VAPA, EIF4E) and FOSL2 (SERBP1, DHX15, MYO1D, VAPA, EIF4E) (Fig. 6C and D).

Identification of FOSL1 binding to COL1A1 and COL1A2 was insightful, since FOSL factors are known to regulate collagen production in other cell types ^81,82^. Additionally, changes in collagen protein levels is correlated with the development of rheumatoid arthritis and osteoarthritis ^75,83^. This suggests a potential involvement of FOSL1 in the incidence of human autoimmunity.

## DISCUSSION

AP-1 factors need to homo- or hetero-dimerize with each other in order to form functional complexes for coordinated gene regulation. AP-1 activity is highly complex, and is systematically regulated at multiple levels, including, the choice of dimerizing partner, post-transcriptional/translational events and interactions with bZIP or other unrelated proteins. Thus, to holistically understand functions of the AP-1 complex, it is critical to investigate the protein-protein interactions of its members.

Our recent functional genomics study showed that FOSL1 and FOSL2 negatively regulate Th17 differentiation in human ^24^. To decipher the mechanisms that govern such functions of FOS-like proteins, we examined their binding partners using a whole-cell proteomics approach. Here, we report the first characterization of FOSL1 and FOSL2 interactomes in human Th17 cells. In addition to their known associations (i.e., c-JUN, JUNB and JUND), we identified many novel binding partners of FOSL proteins. Functional enrichment analysis found a majority of these interactors to be associated with RNA binding activity and translational initiation. RBPs regulate gene-expression by post-translationally modifying stability and splicing of RNA molecules. Although a previous study in cancer cells indicates RBP-mediated regulation of FOS activity ^49^, our findings for the first time holistically reveal a potential cross-talk between these protein groups. Further characterization on this line could broaden the horizons for AP-1 signaling mechanisms in Th17 cells.

Synergistic TFs bind composite regulatory elements through physical interactions between two or more of the candidate factors ^84^. Such mechanisms could be used to integrate distinct signaling pathways and create unified cellular responses. In our analysis, 29 proteins were found to share interactions with FOSL1 and FOSL2. Since both factors alter human Th17 differentiation in a similar fashion, their tendency to bind to common partners suggests functional cooperativity. Intriguingly, the shared hits included RUNX1, SIRT-1 and JUNB, all of which positively regulate Th17 differentiation in mouse ^10,11,65,66,85^. If they have similar roles in the human counterpart, they could antagonize FOSL functions. Interaction-based mechanisms are commonly used by inhibitory proteins to dampen the activity of target TFs ^86,87^. In this respect, our findings suggest that FOSL proteins impair Th17 signaling by binding and sequestering the factors that support the lineage. Remarkably, JUNB and RUNX1 interact with both positive and negative regulators of Th17-fate, owing to which they can perform context-dependent functions. Additional mechanisms, such as post-translational modifications, differential expression profiles and protein stability dynamics, may determine the outcome of their regulatory complexes.

A recent study by He et al. revealed interacting partners of FOSL1 in triple negative breast cancer cells, many of which were reproducibly identified in our analysis (COL1A2, JUN, JUNB, CLTB, CLTC, FUBP3, KHDRBS1, RBM14, DDX17, HNRNPR, and XRN2) ^88^. In addition, we found FOSL1 to associate with clathrin-binding adaptor proteins. This may be attributed to the non-endocytotic roles of clathrin, which involve its translocation to the nucleus to activate gene-transcription ^89^. Furthermore, network analysis highlighted a link between the clathrin binding cluster and double-stranded RNA binding. In relation to this, our study provides insights into the established role of clathrin-mediated endocytosis in the cellular uptake of pathogen-derived double-stranded RNAs ^90,91^. Follow-up experiments are required to determine the actual involvement of RBPs in this process.

FOSL1 was also observed to uniquely interact with factors such as histone deacetylase 2 (HDAC2) and poly-C binding protein 1 (PCBP1), which have reported roles in Th17 regulation. HDAC2 is a global modifier of gene expression that suppresses IL-17 transcription and, thereby, reduces colitis scores ^78^. In contrast, PCBP1 is a ferritin iron regulator that promotes Th17-pathogenicity and autoimmunity ^*52,92*^. These findings indicate that FOSL1 may control the lineage by associating with both activator and repressor complexes. Other novel partners of FOSL1 included SWI/SNF family proteins (SMARCA2, SMARCB1, SMARCC1, SMARCC2, SMARCD2, SMARCE1) and RNA helicase DEAD-box proteins (DDX6, DDX1). Interestingly, several members of these protein families are upregulated upon Th17-initiation ^20,76^, which hints at their involvement in development of the lineage.

The cytoskeleton plays an integral role in transducing extracellular signals to the nucleus ^93^. We found FOSL2 to interact with several proteins involved in actin filament binding (ADD1, ADD3, MYO1B, MYO1D, CAPZA2, ABLIM1, DBNL, CORO1C and FLII). Depolymerization of actin or microtube networks is known to activate c-JUN function, via the JNK/p38 signaling pathway ^94^. c-JUN expression is also induced by actin-binding proteins, such as profilin ^95^. Since JUN emerged as a shared interactor of FOSL1 and FOSL2 in our study, the above findings propose the involvement of cytoskeletal dynamics in regulating FOSL-mediated Th17 networks.

In summary, this study uncovers, for the first time, the global binding partners of FOSL1 and FOSL2 in human T cells, with an emphasis on their shared interactors. Our analysis identified several novel protein-protein associations and molecular functionalities as a part of FOSL-signaling networks. Moreover, the binding of key Th17-regulators to both FOSL1 and FOSL2 highlights the possible mechanisms that mediate the coordinated influence of these factors on the Th17 lineage. It is established that PPI networks of TFs are significantly altered in cases of mutations or disease ^96^. Since FOS-like proteins have important implications in the development of autoimmune disorders ^97-101^, their interactomes could serve as a crucial resource in the field of disease-biology. Studying the changes in their protein-protein interactions under adverse physiological conditions, could help predict diagnostic markers and therapeutic targets for Th17-associated pathologies.

## ACKNOWLEDGMENTS

We thank all voluntary blood donors and personnel of Turku University Hospital, Department of Obstetrics and Gynecology, Maternity Ward (Hospital District of Southwest Finland) for the umbilical cord blood collection. We are grateful to Marjo Hakkarainen and Sarita Heinonen for their excellent technical help. We duly acknowledge Turku Proteomics Facility supported by Biocenter Finland, for their assistance. The Finnish Centre for Scientific Computing (CSC) and and ELIXIR Finland are acknowledged for computational resources.

A.S was supported by Erasmus Mundus Scholarship, University of Turku (UTU) and Council of Scientific & Industrial Research (CSIR), Government of India. S.K.T. was supported by the Juvenile Diabetes Research Foundation Ltd (JDRF; grant 3-PDF-2018-574-A-N); SG received grants from the Centre of Excellence in Epigenetics program (Phase II) of the Department of Biotechnology (BT/COE/34/SP17426/2016), Government of India, the JC Bose Fellowship (JCB/2019/000013) from the Science and Engineering Research Board, Government of India, and the Academy of Finland (AoF/315585). R.L. received funding from the Academy of Finland (grants 292335, 292482, 298732, 294337, 298998, 31444, 315585, 319280, 329277, 323310, 331790) by grants from the JDRF, the Sigrid Jusélius Foundation (SJF); Jane and Aatos Erkko Foundation, The Finnish Diabetes Foundation, The Novo Nordisk Foundation and the Finnish Cancer Foundation. Our research is also supported by InFLAMES Flagship Programme of the Academy of Finland (decision number: 337530) and University of Turku Graduate School (UTUGS).

## AUTHOR CONTRIBUTIONS

A.S designed and performed the experiments, analysed data, prepared figures, and wrote the manuscript; S.D.B. designed and performed experiments, analysed the proteomics data, prepared figures, and wrote parts of the manuscript; S.K.T. initiated the study, designed and performed the experiments, provided expertise and wrote parts of the manuscript, T.B performed experiments and prepared figures; R.B. prepared cultures and assisted with experiments; O.R. and R.M provided expertise and guidance, and edited the manuscript; S.G. supervised, provided expertise, and edited the manuscript; R.L. designed the study setup, provided expertise, participated in the interpretation of the results, provided guidance and supervision, and wrote the manuscript. All authors have contributed to the manuscript.

## SUPPLEMENTARY INFORMATION

### Supplementary Figures and Legends

**Fig. S1.**
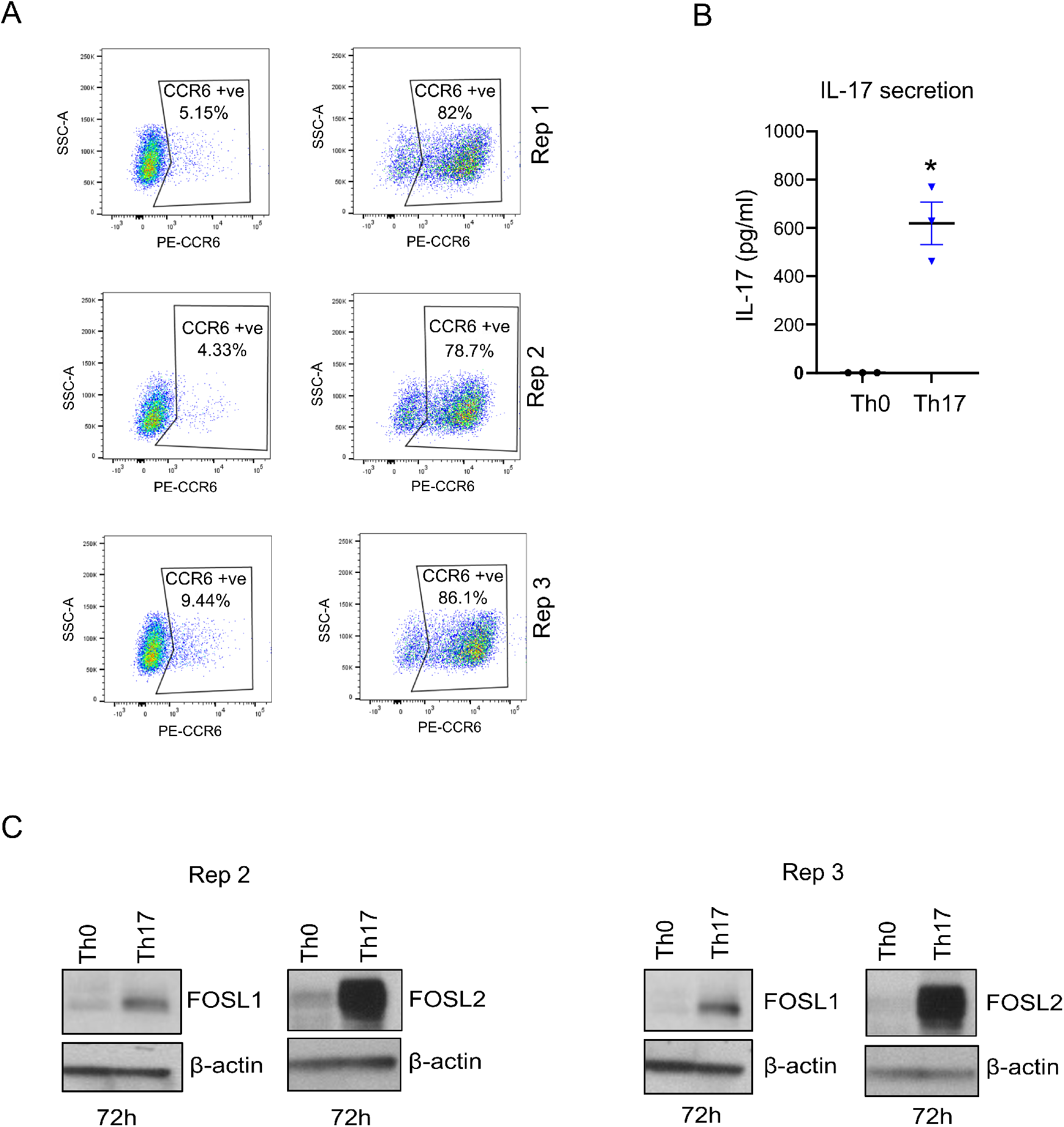
Th17-differentiation and associated expression of FOSL1 and FOSL2. **A**. Flow cytometry plots show percentage of CCR6 positive cells in Th0 and Th17 cultures at 72h of polarization. Cultures with polarization efficiencies equivalent to the ones shown here were used for proteomic analysis. Data for three biological replicates is shown. **B**. Graph shows ELISA values for IL-17 secretion in 72h cultured Th0 and Th17 cells. IL-17 values were first normalized to live cell count and then to Th0 control. Data is based on three biological replicates. Two-tailed students t-test was used to determine statistical significance (*p < 0.05). **C**. Immunoblots show protein levels of FOSL1 (left) and FOSL2 (right) in naive CD4^+^ T cells cultured under activating (Th0) or Th17-polarizing conditions for 72h. Actin was used as loading control. Blots represent biological replicates for Fig. 1A.

**Fig. S2.**
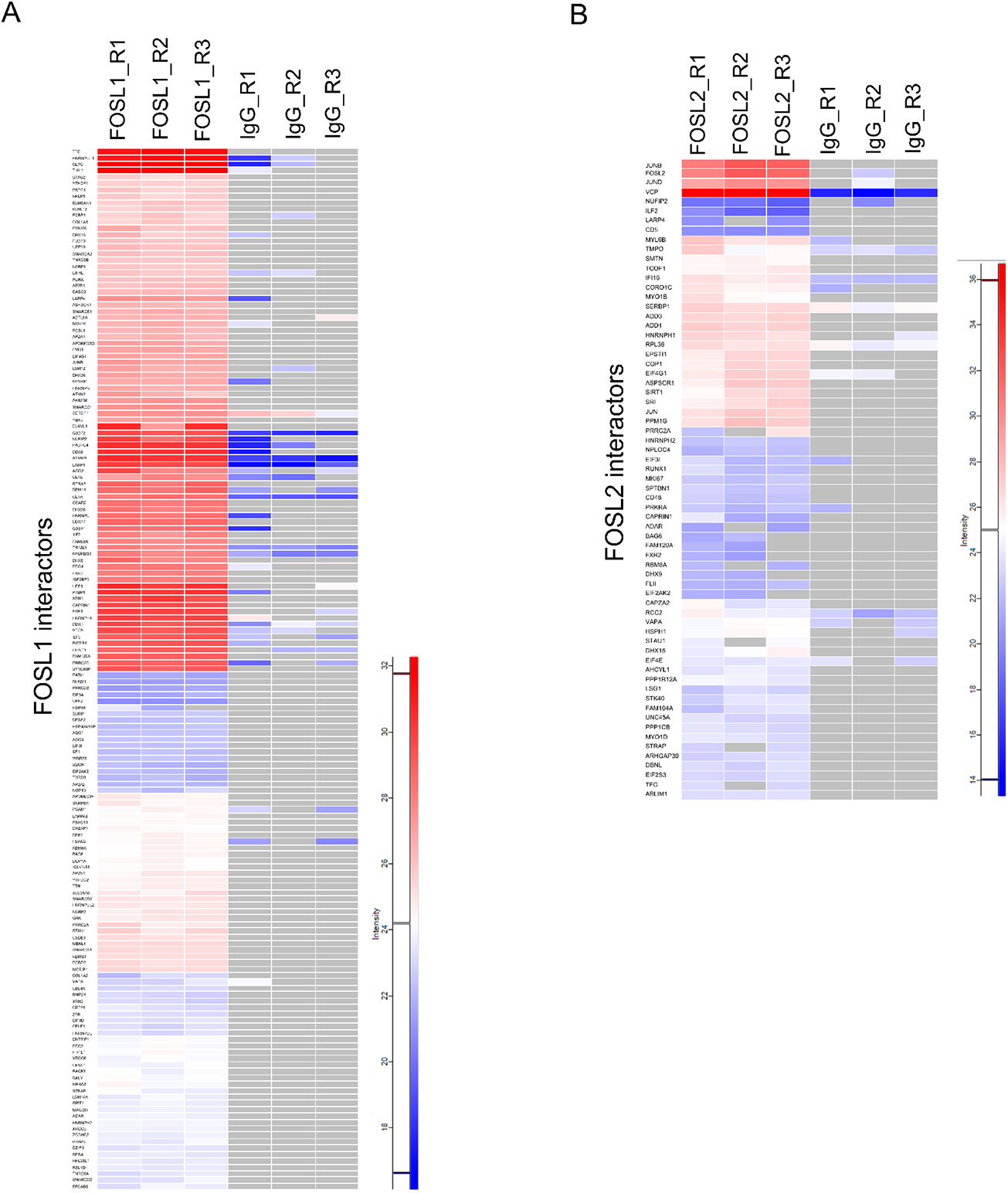
FOSL1 and FOSL2 interacting proteins in human Th17 cells. **A & B**. Heatmaps shows Log_2_intensity values for the FOSL1 (panel A) and FOSL2 (panel B) interactors that were identified by MS analysis in Th17 cells. Grey color represents undetected or missing proteins.

**Fig. S3.**
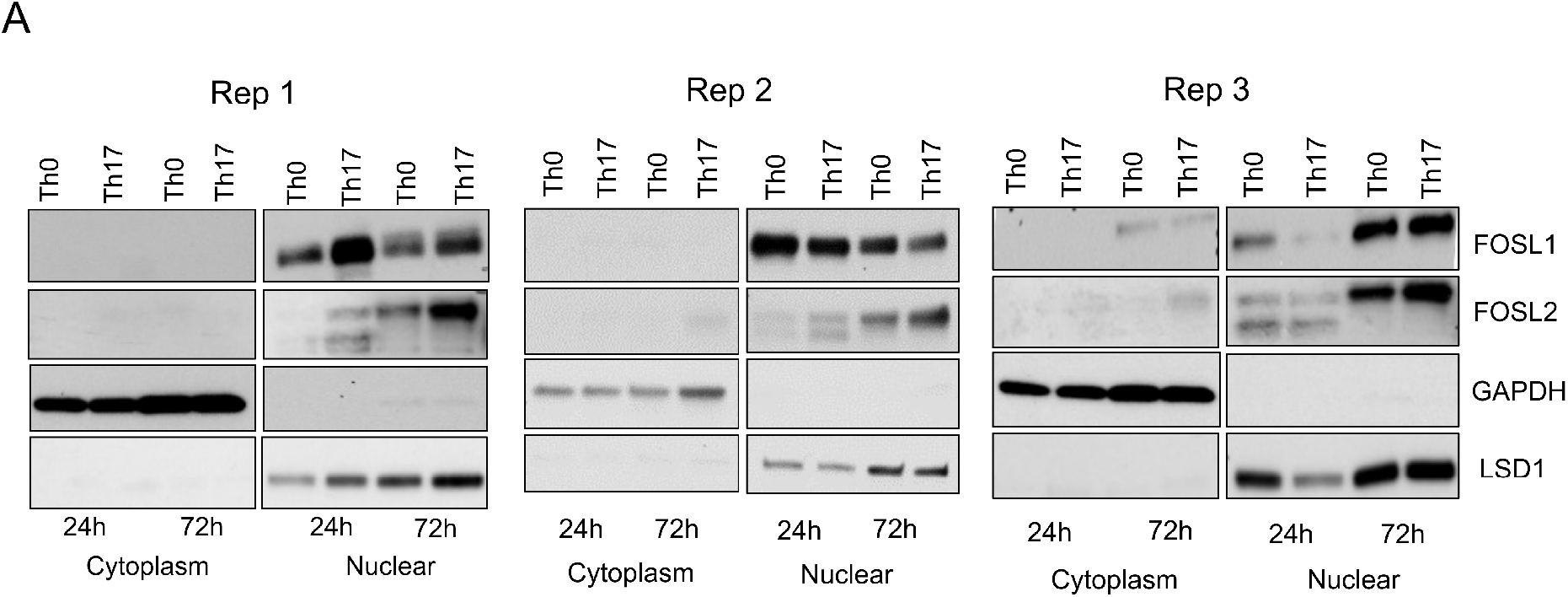
Subcellular fractionation of FOSL1 and FOSL2 in human Th17 cells. **A**. Th0 and Th17 cell lysates (24h and 72h) were fractionated and further probed for expression of FOSL1 and FOSL2 using western blotting. GAPDH and LSD1 mark cytoplasmic and nuclear fractions, respectively. Immunoblots show biological replicates for Fig. 3A.

**Fig. S4.**
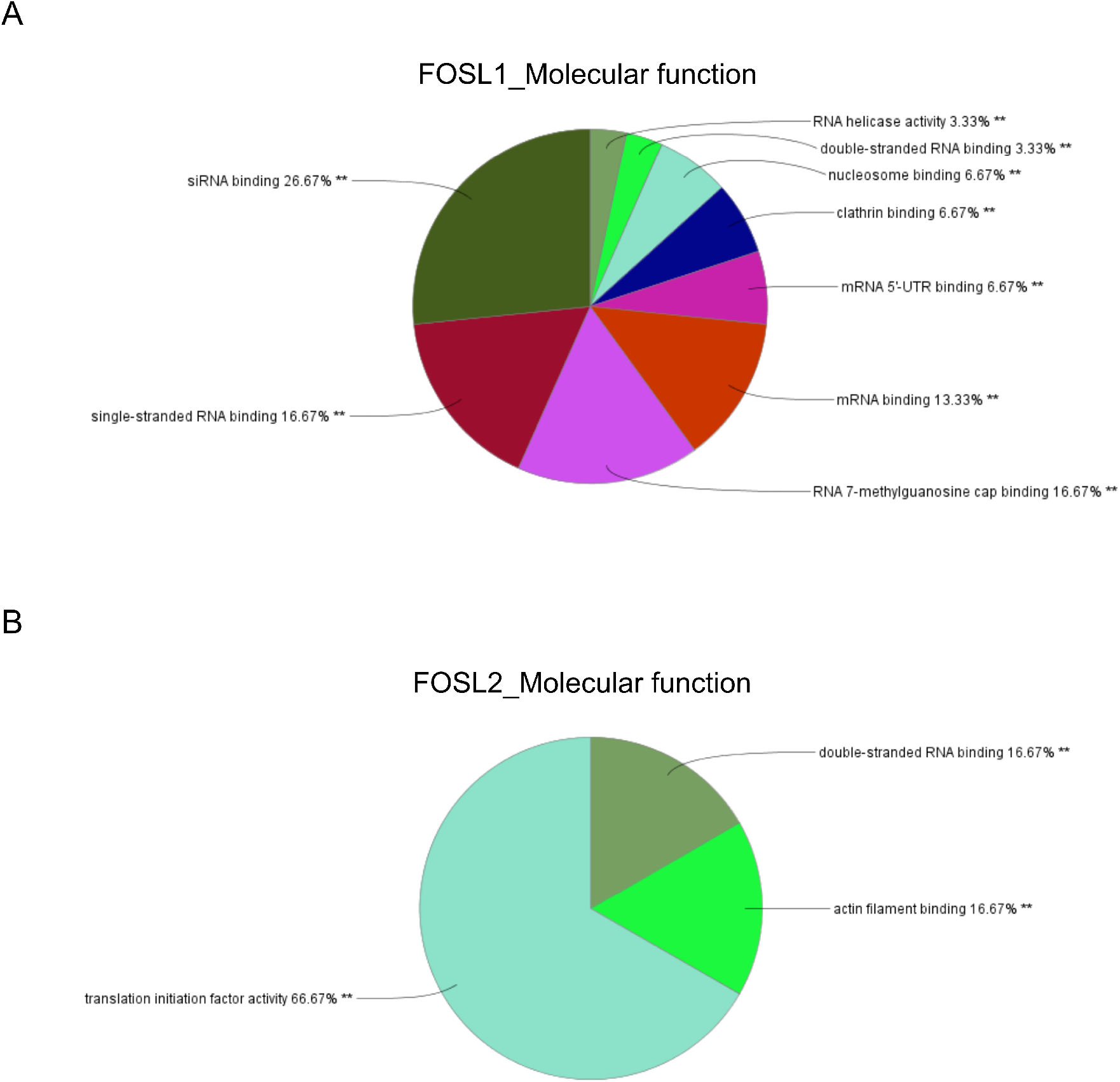
Molecular functionalities of FOSL1 and FOSL2 interactors. **A & B**. Pie charts illustrate the classification of FOSL1 (panel A) and FOSL2 (panel B) interacting partners on the basis of their molecular function. ClueGO and CluePedia plugins from Cytoscape was used to create the charts.

**Fig. S5.**
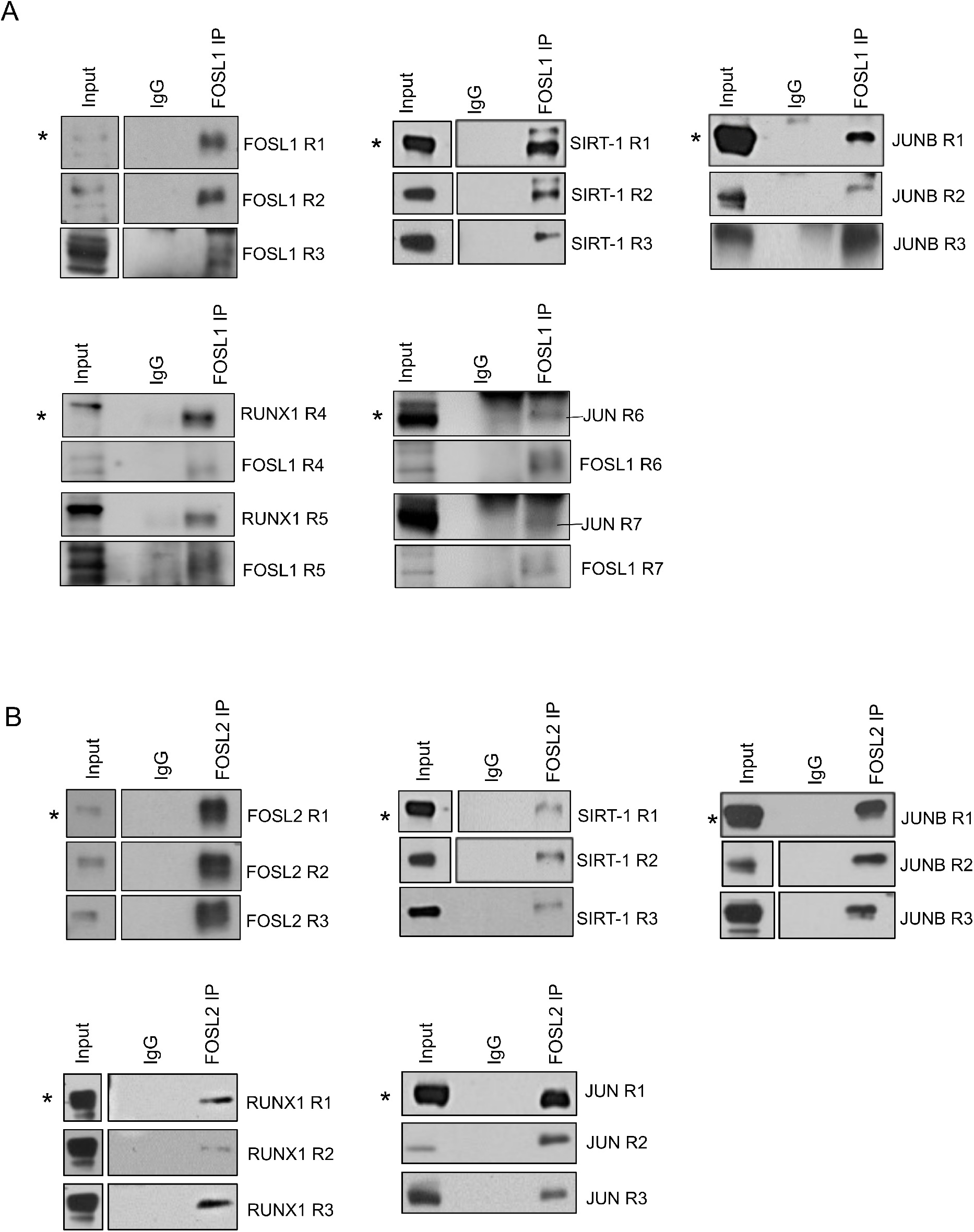
Validation of shared interactors of FOSL1 and FOSL2 using immunoblot analysis. **A & B**. FOSL1 (panel A) or FOSL2 (panel B) protein was immunoprecipitated from 72h cultured Th17 cells and immunoblotting was performed to confirm their shared interactions with SIRT-1, JUNB, RUNX1 and JUN. Blots show lanes for total lysate (input), control IgG IP and FOSL1/FOSL2 IP. R1-R7 in panel A and R1-R3 in panel B represent different biological replicates for the FOSL1 and FOSL2 IP blots in Figs. 6A and 6B, respectively (the representative blots shown in the main figures are denoted here with *).

